# Integrated effect of plant growth regulators with boron sources on some biological parameters of sugar beet

**DOI:** 10.1101/839068

**Authors:** Marwa A. Qotob, Mostafa A. Nasef, Heba K. A. Elhakim, Olfat G. Shaker, Nader R. Habashy, Ismail A. Abdelhamid

**Affiliations:** Soils, Water and Environment Research Institute, Agriculture Research Center, Giza, Egypt; Biochemistry department, Faculty of Science, Cairo University, Giza 12613, Egypt; Department of Biochemistry, Faculty of Medicine, Cairo University, Egypt; Department of Chemistry, Faculty of Science, Cairo University, Giza12613, Egypt

**Keywords:** Proline, GA_3_, Boric acid, B-NPs, Macro-nutrient fertilizers, Sucrose quality, sugar beet, Betaine, Choline

## Abstract

Any improvement in agricultural systems that results higher production aimed to reduce negative environmental impacts and enhance sustainability plant growth regulators (PGRs) such as gibberellin have similar physiological and biological effects to those of plant hormones, and therefore used widely in agriculture to minimize unwanted shoot growth without lowering plant productivity.

An experimental field was conducted at Giza Experimental Station Egypt, on sugar beet plants (*Beta vulgaris* L. *var.* Sara poly) with some plant growth regulators (gibberellin and proline) foliar application at three rates of zero (control), 100 and 200 mg l^−1^ and boron sources (Boric acid and B-NPs) with 75% of macro-nutrients from full dose.

The main target of this study to evaluate another plant growth regulator source like proline which is safer than gibberellin for maximizing sugar beet biological parameters to reduce the gap between sugar consumption and production in presence of boron sources.

Data showed that the foliar applications of gibberellin (GA_3_) at rate 100 mg l^−1^ and proline at 200 mg l^−1^ were found to be the more effective without significant differences for plant growth, productivity and quality may be due to increased N use efficiency, especially at sub-optimal macro nutrient fertilizers. Regard to boron sources, B-NPs had positive effect on all biological parameters under study due to sugar transport, cell membrane synthesis, nitrogen fixation, respiration, carbohydrate metabolisms, root growth, functional characteristics and development.

## 1. Introduction

Sugar beet (the raw material of the beet sugar factory) composition is important to both the sugar beet farmer and the factory. Sugar (sucrose) and non-sugar (non-sucrose) content determine the quality of the sugar beet where, high sugar and low non-sugar content is desirable. So it is important to evaluate the chemical quality of sugar beet roots in order to evaluate their quality for sugar production. Root yield and technical quality of sugar beet are strongly influenced by weather conditions. The technical quality of sugar beet is essential for economical sugar manufacturing (Asadi, 2006). The wide majority of beets are grown by independent farmers, who are contracted by the factories directly. The industrial demand for sugar beets is increasing, which provides a higher price, incentivizing many farmers to plant more beets. FAS (Foreign Agricultural services) Cairo is increasing sugar beet area harvested in MY 2019/20 to 250,000 ha. Post is revising down last year’s harvested area to 225,000 hectares instead of 230,000. With the decreased area harvested, post also revised down MY 2018/19 production to 9.5 MMT, six percent below earlier estimates. Macro-nutrients fertilization is among the vital factors affecting growth, quality and productivity of sugar beet, thus application the suitable rates of nitrogen, phosphorus and potassium one of the favorable factors for increasing sugar beet productivity and quality. The proper fertilization reference can be given only based on the soil fertility. It must be determining optimum nitrogen rate, which produce the maximum root, sugar yields and quality parameters (Seadh, 2012).

Plant growth regulators are hormones that widely used in agriculture to increase plant growth and reproduction. The commonly used class of plant growth regulators includes plant growth hormones such as cytokinin, and gibberellic acid (Fukao and Bailey-Serres, 2008).

Gibberellins involved in a number of cellular processes that regulate seed germination and growth of aerial plant parts, including floral induction and fruit development (Spaepen, S.; Vanderleyden, J. and Okon, 2009). The effect of spray of gibberellic acid (GA_3_) at very low concentrations could be exploited beneficially as its natural occurrence in plants in minute quantities is known to control their development. It is an established phytohormone used commercially for improving the productivity and quality of a number of crop plants (King and Evans, 2003).

Proline is the most widely distributed metabolite that accumulates under stress conditions (Delauney and Verma, 1993) the significance of this accumulation in osmotic adjustment in plants is still debated and varies from species to species (Hoai et al., 2003). The crystallization of sugar in the industrial processing of beet root in sugar refineries may be jeopardized by accumulation of compounds, such as proline and glucose, because they lead to the formation of coloured components that reduce the quality of beet roots (Monreal et al., 2007).

Boron is one of the important micronutrient among essential elements for plant growth, and plays a significant role in the physiological and biochemical processes within plants (Tariq and Mott, 2006). Boron plays a key role in higher plants by facilitating the short-and long-distance transport of sugar via the formation of borate-sugar complexes. In addition, boron may be of importance for maintaining the structural integrity of plasma plant cells membranes. This function is likely related to stabilization of cell membranes by boron association with some membrane constituents (Brown et al., 2002).

Nanotechnology to precisely detects and delivers the correct quantity of nutrients and pesticides and increases the bioavailability (Goudar et al., 2018) which promote productivity while ensuring environmental safety and higher use efficiency. Nano boron has many merits like quick and easy uptake by plants. It has lower tendency to leach via soil and appear its impact for shorter times. It improves solubility and dispersion of insoluble nutrients in soil, reduces soil fixation.

The main target of this study using some plant growth regulators *i.e.* gibberellin and proline to increase sugar beet productivity and its biological parameters without hazard effect on human beings and environment in presence of boron sources (Boric acid and Nano boron) with controlled use of macro-nutrient fertilizers.

## 2. Materials and methods

A field experiment was conducted on a clay texture soil at El Giza Agricultural Research Station, Egypt (located between 30° N, 31°: 28 E at an altitude 19 meters above sea level) and cultivated with sugar beet (*Beta vulgaris var.* Sara poly) during winter season of 2018. The current study aimed to identify the direct beneficial effects on applying some plant growth regulators as gibberellin and proline at different rates (100, 200 mg l^−1^) with two boron sources (boric acid and nano boron, B-NPs) at recommended dose for sugar beet (0.48 Kg B acre^−1^) at full dose and 75% from full dose of macro-nutrient fertilizers. The nano boron (B-NPs) analysis of X-ray pattern in (Fig. 1).

**Fig. 1.**
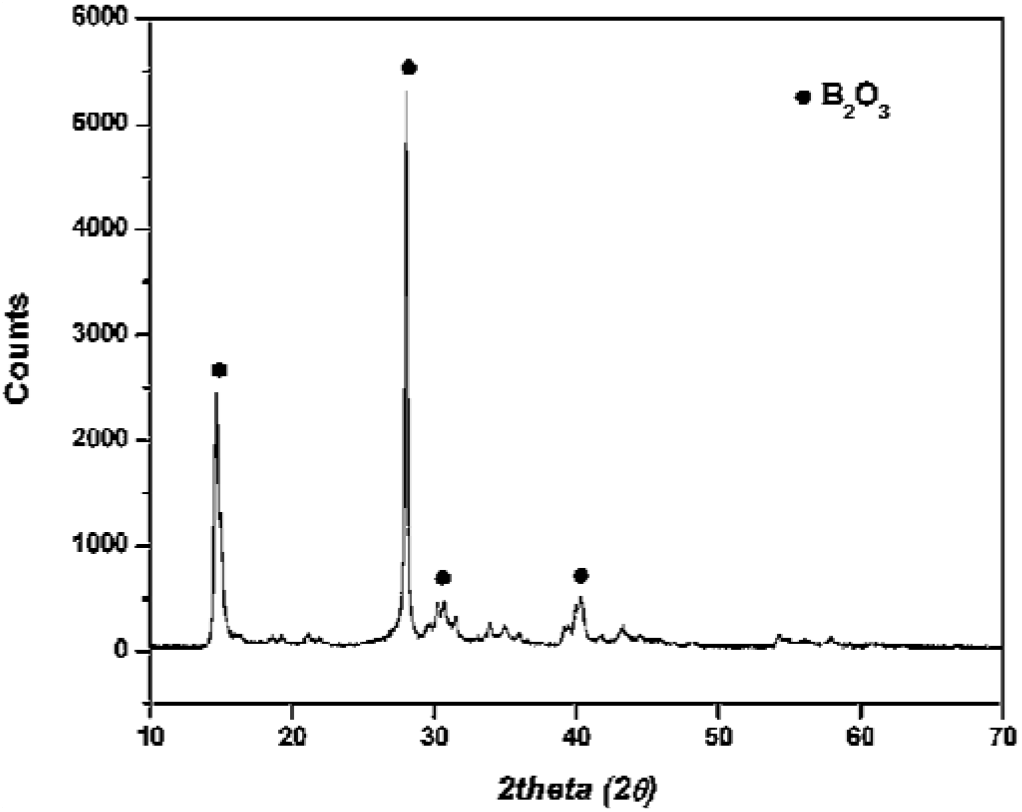
X-ray pattern of the B_2_O_3_ NPs (1 – 45 nm)

The experiment was laid-out in split- split plot design with three replicates as follows:

a. The main plots were 75% from full dose. Calcium super phosphate (15.5% P_2_O_5_) was added at rate 112.5 P_2_O_5_ Kg acre^−1^ during the soil preparation. Nitrogen was applied at rate of 56.5 Kg N acre^−1^ as urea (46.5% N) in three equal doses after 21, 45 and 60 days from planting. Potassium sulphate (50% K_2_O) at rate of 37.5 Kg K_2_O acre^−1^ was added in two equal doses after 30 and 50 days from planting. Foliar applications of both plant growth regulators and boron were applied after 45 and 60 days from sowing. All cultural practices for growing sugar beet were done as recommended.
b. The sub plots were plant growth regulators *i.e.* GA_3_ (Natural Enterprise Co.) and proline (Alfa Aesar Co.) at three rates as a foliar application (Zero, 100 and 200 mg l^−1^).
c. The sub – sub plots were applied foliar with two boron sources boric acid (Aldrich Co.) and B-NPs (Yara Fertiliser Co.) at the recommended dose for sugar beet (0.48 Kg B acre^−1^)

An agricultural soil sample (0–30 cm depth) was used for the study. It was air-dried, ground and sieved with a 2 mm sieve. Some of its properties were estimated according to (Page et al., 1982). The total N was determined by distillation in a Macro-Kjeldahl (Gerhardt model VAP 30 S). Total P was estimated colorimetrically using stannous chloride mixture and measured by UV/Vis spectrophotometer (JENWAY model 6705 UV/Vis), while K^+^ and Na^+^ concentrations were measured by flame photometer (JENWAY model PFP7). The concentrations of Ca^+2^, Mg^+2^, Fe, Mn, Zn, Cu and B were measured by ICP-AAS spectrophotometer (Agilent Technologies model 8800) (Jackson, 1959), (Cottenie et al., 1982), Tabulated in (Table 1).

**Table 1.** Chemical and physical characteristics of initial soil under investigation.

| Soil characteristics | Value | Soil characteristics | Value |
| --- | --- | --- | --- |
| Particle size distribution%: |  | Soluble cations (soil paste, mmol<sub>c</sub>l<sup>-1</sup>): |  |
| Sand | 26.2 | Ca<sup>+2</sup> | 2.35 |
| Silt | 29.3 | Mg<sup>+2</sup> | 1.20 |
| Clay | 44.5 | Na<sup>+</sup> | 6.85 |
| Soil textural class | Clay* | K<sup>+</sup> | 5.13 |
| Soil chemical properties: |  | Soluble anions (soil paste, mmol<sub>c</sub>l<sup>-1</sup>): |  |
| pH (1:2.5 soil water suspension) | 8.95 | CO<sub>3</sub><sup>-2</sup> | 0.00 |
| CaCO<sub>3</sub> % | 4.82 | HCO<sup>-3</sup> | 3.15 |
| Organic matter % | 1.53 | Cl<sup>-</sup> | 6.40 |
| EC<sub>e</sub> (dS m<sup>-1</sup>, soil past) | 1.69 | SO<sub>4</sub><sup>-2</sup> | 5.95 |
| Soil physical properties: |  | Available macro- and micronutrients mg kg<sup>-1</sup> |  |
| Bulk density, g cm<sup>-3</sup> | 1.20 | N | 46.34 |
| Sodium adsorption ratio (SAR) | 5.15 | P | 18.56 |
| Exchangeable sodium (ESP) | 4.50 | K | 349.1 |
| Saturation (SP) | 70.3 | Fe | 41.2 |
| CEC** cmol<sub>c</sub> kg<sup>-1</sup> | 53.2 | Mn | 26.1 |
| Moisture content %: |  | Zn | 2.43 |
| Field capacity | 27 | B | 1.13 |
| Wilting point | 16 |  |  |
| Available water | 11 |  |  |

\* Using USDA Soil Texture Triangle, after (Twarakavi et al., 2010).
\*\* CEC= Cation exchange capacity

Chemical analyses of sugar beet plants were carried out on the samples to determine boron by ICP (Inductively coupled plasma) spectrometry (model, Ultima 2 JY Plasma) (Jackson, 1959), (Cottenie et al., 1982), total sugar were determined in sugar beet roots calorimetrically with the picric acid method as described by (Thomas and Dutcher, 1924), betaine was determined by (Focht et al., 1956), choline was determined according to (Gimesi and Szász, 1974), proline was investigated by (Carillo and Gibon, 2011) by the absorbance of the extract measured Spectrophotometer using by JENWAY 6705 UV/Vis., gibberellin was determined using gas spectroscopy model Trace 1310 GC as described by (Moritz and Monteiro, 1994), total soluble solids was determined by (Mariani et al., 2014) using refractometer (model J57HA), determination of α- Amino nitrogen by (Shtanheev et al., 1998), purity was measured by (Guyot, 1967).

Data statistically analyzed by the analysis of variance (ANOVA) was carried out to determine the statistical significance using the least significant difference at level at 0.05 (Gomez and Gomez, 1984).

## 3. Results and discussion

The aim of this study to reduce the consumption of mineral fertilizers through using plant growth regulators (gibberellin or proline) at the same time evaluate possibility replace the gibberellin by proline which showed its harmful effect on animal and human cells according to (Wang et al., 2011) who stated that the present study clearly demonstrated that PGRs including gibberellin and cytokinin may cause acute toxicity and teratogenic effect in both neonate and embryo cells.

### 3.1. Pigment content in sugar beet

The obtained data in Table 2 revealed that the foliar application of plant growth regulators (gibberellin and proline) increased the pigment content *i.e*. chlorophyll-a; chlorophyll-b and carotenoids where the mean values at rate 100 mg l^−1^ gibberellin and 200 mg l^−1^ proline were 6.15 and 5.44 g100 g^−1^ FW for chlorophyll-a, 3.39 and 2.99 g100 g^−1^ FW for chlorophyll-b, 0.916 and 0.887 g100 g^−1^ FW, respectively. Exogenous application of gibberellins (GAs) has shown enhanced activities of carbonic anhydrase, nitrate reductase (Afroz, S., Mohammad, F., Hayat, S., Siddiqui, 2005), CO_2_ fixation, stomatal conductance (Bishnoi and Krishnamoorthy, 1991) and ribulose-1,5-biphosphate carboxylase/oxygenase (Yuan and Xu, 2001). GAs alters membrane permeability to ions (Gilroy, S. and Jones, 1992) and improves translocation potential to the sink (Peretó and Beltrán, 1987). The physiological and biochemical modes of action of GA_3_ is likely responsible for increasing shoot dry biomass plants. Generally, lower concentrations of plant growth regulators are biochemically more active with higher concentrations becoming toxic (Kulkarni et al., 2013). Proline, a multifunctional amino acid, besides acting as an excellent osmolyte is also known for stabilizing subcellular structures such as proteins and cell membranes, scavenging free radicals, balancing cellular homeostasis and signaling events and buffering redox potential under stress conditions (Hayat et al., 2012). It could be reflecting on maintaining the nutrient status in roots.

**Table 2.** Integrated effect of plant growth regulators and mineral fertilizer rates with different boron sources on sugar beet pigments.

| Boron sources | Gibberellin (mg l<sup>-1</sup>) |  |  |  | Proline (mg l<sup>-1</sup>) |  |  |
| --- | --- | --- | --- | --- | --- | --- | --- |
|  | Control | 100 | 200 | Mean | 100 | 200 | Mean |
| **Chlorophyll-a (g 100g<sup>-1</sup> FW)** |  |  |  |  |  |  |  |
| Control | 3.14 | 5.59 | 4.99 | 4.57 | 4.10 | 5.12 | 4.12 |
| Boric acid | 3.23 | 6.18 | 5.49 | 4.96 | 4.83 | 5.25 | 4.44 |
| Nano B | 3.46 | 6.67 | 5.70 | 5.28 | 4.90 | 5.94 | 4.77 |
| Mean | 3.28 | 6.15 | 5.39 | 4.94 | 4.61 | 5.44 | 4.44 |
| **Chlorophyll-b (g 100g<sup>-1</sup> FW)** |  |  |  |  |  |  |  |
| Control | 1.33 | 2.71 | 2.68 | 2.24 | 2.45 | 2.70 | 2.16 |
| Boric acid | 1.59 | 3.65 | 3.22 | 2.82 | 2.60 | 2.92 | 2.37 |
| Nano B | 1.72 | 3.80 | 3.50 | 3.01 | 2.73 | 3.35 | 2.60 |
| Mean | 1.55 | 3.39 | 3.13 | 2.69 | 2.59 | 2.99 | 2.38 |
| **Carotenoids (g 100g<sup>-1</sup> FW)** |  |  |  |  |  |  |  |
| Control | 0.673 | 0.831 | 0.728 | 0.744 | 0.731 | 0.802 | 0.735 |
| Boric acid | 0.685 | 0.937 | 0.746 | 0.789 | 0.769 | 0.892 | 0.782 |
| Nano B | 0.708 | 0.981 | 0.771 | 0.820 | 0.824 | 0.978 | 0.837 |
| Mean | 0.689 | 0.916 | 0.748 | 0.784 | 0.775 | 0.881 | 0.783 |
| **L.S.D. at 0.05** |  |  |  |  |  |  |  |
|  | B | G | R | B x G | B x R | G x R | B x G x R |
| Chlorophyll-a | 0.61 | 1.01 | 0.55 | 0.57 | 0.56 | 0.70 | 0.50 |
| Chlorophyll-b | 0.37 | 0.85 | 0.45 | 0.51 | 0.11 | 0.30 | 0.35 |
| Carotenoids | 0.02 | 0.03 | 0.10 | 0.09 | 0.04 | 0.15 | 0.02 |

B = Boron sources; G = Plant growth regulator sources and R = Rate of Plant growth regulator

This report is in conformity with the increased nitrate content of roots by exogenous application of proline (Alyemeni et al., 2016). exogenous application of proline mitigated the decreased plant growth caused by stress is through increasing antioxidant system, relieving oxidative damage, improving the synthesis of compatible solutes, and accelerating proline accumulation, which reflected on enhancing photosynthesis (Anjum et al., 2011). Also, the chlorophyll molecules are the membrane bound structures whose stability depends highly on the integrity of the membrane structure which is possibly maintained by proline as it acts as a membrane stabilizer (Ashraf, M. and Foolad, 2007).

Regard to boron sources, data showed that foliar application of nano boron on sugar beet plants was more respond than boric acid where the chlorophyll-a, chlorophyll-b and carotenoids contents were 3.13, 1.58 and 0.695 g100 g^−1^ FW for nano boron and 3.06, 1.40 and 0.662 g100 g^−1^ FW for boric acid, respectively. This result was in agreement with (Hassan et al., 2013) who reported that application of boron increase net photosynthetic rate which may be attributed to the increase in chlorophylls content of leaves. Furthermore, application of boron increased the activity of catalase and glutathione reductase, which act as antioxidants thus saving the electron transport mechanism of plant from getting oxidized by free radicals like superoxide radicals, singlet oxygen radicals (Wojcik et al., 2008). Also, (Davarpanah et al., 2016) indicated that the foliar application of nano-B fertilizers in plant increased the leaf concentrations of both microelements, reflecting the improvements in nutrient status.

In general, it could be noted that the interaction between foliar application of plant growth regulators accompanied with boron sources used give more respond than plant growth regulators or boron sources sole and the more effective treatment were 100 mg l^−1^ gibberellin and 200 mg l^−1^ proline with nano boron without any significant differences.

### 3.2. Residual content of applied plant growth regulators and boron sources used in sugar beet roots

Data showed in Table 3 that the foliar application of plant growth regulators (Gibberellin or Proline) at two rates 100 and 200 mg kg^−1^ on sugar beet roots residual content increased by application of gibberellin. The obtained data was in agreement with (Kim et al., 2008) who reported that GA_3_ applications significantly increased endogenous GA_3_ content in the plants. On contrast, proline application did not affect the residual content of gibberellin when applied at both rate compared to control treatment. According to (Osman, 2015) exogenous application of proline might be not only accelerated the translocation process of amino acids from source to sink, but also suppressed the conversion process from amino acids to proteins.

**Table 3.** Residual content of plant growth regulators applied with different boron sources on sugar beet roots.

| Boron sources (B) | Gibberellin (mg l<sup>-1</sup>) |  |  |  | Proline (mg l<sup>-1</sup>) |  |  |
| --- | --- | --- | --- | --- | --- | --- | --- |
|  | Control | 100 | 200 | Mean | 100 | 200 | Mean |
| Proline (μmol g<sup>-1</sup> FW) |  |  |  |  |  |  |  |
| Control | 2.16 | 2.39 | 2.16 | 2.28 | 2.25 | 2.45 | 2.33 |
| Boric acid | 2.29 | 3.06 | 2.74 | 2.75 | 2.69 | 2.78 | 2.64 |
| Nano B | 2.41 | 3.11 | 2.85 | 2.86 | 2.84 | 3.01 | 2.82 |
| Mean | 2.45 | 2.85 | 2.58 | 2.63 | 2.59 | 2.75 | 2.60 |
| Gibberellin (mg kg<sup>-1</sup>) |  |  |  |  |  |  |  |
| Control | 13.05 | 15.68 | 14.70 | 14.48 | 13.46 | 13.61 | 13.37 |
| Boric acid | 13.07 | 16.25 | 15.54 | 14.95 | 13.62 | 13.89 | 13.53 |
| Nano B | 13.15 | 16.75 | 15.80 | 15.23 | 13.80 | 14.15 | 13.70 |
| Mean | 13.09 | 16.23 | 15.35 | 14.89 | 13.63 | 13.88 | 13.53 |
| Boron (mg kg<sup>-1</sup>) |  |  |  |  |  |  |  |
| Control | 46.21 | 65.82 | 60.45 | 56.97 | 61.09 | 67.38 | 57.71 |
| Boric acid | 49.82 | 78.64 | 66.12 | 63.58 | 65.62 | 70.53 | 60.71 |
| Nano B | 52.45 | 84.35 | 72.30 | 68.80 | 73.86 | 78.81 | 67.47 |
| Mean | 49.49 | 76.27 | 66.29 | 63.12 | 66.86 | 72.24 | 61.96 |
| L.S.D. 0.05 |  |  |  |  |  |  |  |
|  | B | G | R | B x G | B x R | G x R | B x G x R |
| Proline | 0.08 | 0.15 | 0.11 | 0.10 | 0.11 | 0.12 | 0.05 |
| Gibberellin | 0.15 | 1.20 | 0.50 | 0.21 | 0.11 | 0.20 | 1.40 |
| Boron | 2.01 | 3.10 | 4.71 | 4.55 | 5.09 | 4.85 | 1.25 |

B = Boron sources; G = Plant growth regulator sources and R = Rate of Plant growth regulator

Also, data showed that the residual content of proline and boron increased by application rates of gibberellin, this may be due to that the gibberellic acid may stimulated plant on building protein and the optimum transform of nutrients although the unavailability of moisture in the plant to a certain level (Al-Shaheen and Soh, 2016). Regard to the application of proline at 100 and 200 mg kg^−1^ proline and boron content were increased by increasing proline concentration. According to (Kahlaoui et al., 2014) the exogenous application of proline leads to a significant increase in the proline accumulation of both organs (leaves and roots) in plant.

Further study showed that amino acids such as proline have a chelating effect on micronutrients when applied together; the absorption and transportation of micronutrients inside the plant is easier, this effect is due to the chelating action, the effect of cell membrane permeability and low molecular weight (Westwood, 1993).

Regard to the foliar application of boron on plants of sugar beet (*Beta vulgaris* L.) in the form of boric acid or B-NPs, the obtained data showed that the mean values of proline content were 2.45, 2.17, 2.62, and 2.35 μmol g^−1^ FW, gibberellin content 13.07, 13.02, 13.15 and 13.09 mg kg^−1^ and boron content 49.82, 45.98, 52.45 and 49.75 mg kg^−1^, respectively. According to (Nilanjan, 2013) stated that compared to the conventional boric acid or Borax fertilizers, all of which are on the macro scale (on the order of micrometers) (macro boric acid”, the boron nanofertilizers of embodiments herein (on the order of nanometers) shows a sharp increase in crop yield (increased biomass, potato tuber yield, and plant weight) and crop quality (less reducing sugar and increased starch content).

Finally, the interaction analysis for 75% NPK from the recommended dose accompanied with foliar application of gibberellin at rate 100 mg l^−1^ and nano boron (NPs) was the most effective treatment than the same interaction with boric acid.

### 3.3. Sugar extraction and quality parameters

Data obtained in (Table 4) revealed that the foliar application of plant growth regulator as a gibberellin at rate 100 mg l^−1^ give the highest values for sucrose, purity and total soluble solids whereby increase the rate of gibberellin at rate of 200 mg l^−1^ the was observed that decreased in all pervious parameters. In contrast, the results showed that the foliar application of proline gave higher response at 200 mg l^−1^ than 100 mg l^−1^ as application rates where, proline, a multifunctional amino acid, besides acting as an excellent osmolyte is also known for stabilizing subcellular structures such as proteins and cell membranes, scavenging free radicals, balancing cellular homeostasis and signaling events and buffering redox potential under stress conditions (Hayat et al., 2012).

**Table 4.** Integrated effect of plant growth regulators and macro-nutrients fertilizer rates with different boron sources on Sugar content and quality.

| Boron sources (B) | Gibberellin (mg l<sup>-1</sup>) |  |  |  | Proline (mg l<sup>-1</sup>) |  |  |
| --- | --- | --- | --- | --- | --- | --- | --- |
|  | Control | 100 | 200 | Mean | 100 | 200 | Mean |
| **Sucrose (%)** |  |  |  |  |  |  |  |
| Control | 16.09 | 17.65 | 16.81 | 16.82 | 16.95 | 17.48 | 16.84 |
| Boric acid | 16.48 | 18.30 | 16.90 | 17.23 | 17.10 | 17.70 | 17.09 |
| Nano boron | 16.69 | 18.90 | 17.60 | 17.79 | 17.84 | 18.65 | 17.73 |
| Mean | 16.42 | 18.28 | 17.10 | 17.24 | 17.30 | 17.94 | 17.22 |
| **Purity (%)** |  |  |  |  |  |  |  |
| Control | 80.42 | 82.80 | 81.89 | 81.60 | 82.07 | 82.43 | 81.54 |
| Boric acid | 80.47 | 83.57 | 82.21 | 82.01 | 82.71 | 82.75 | 81.90 |
| Nano boron | 80.62 | 84.51 | 82.85 | 82.56 | 82.80 | 83.99 | 82.47 |
| Mean | 80.50 | 83.63 | 82.32 | 82.01 | 82.53 | 83.06 | 81.97 |
| **α- Amino nitrogen (%)** |  |  |  |  |  |  |  |
| Control | 1.24 | 1.12 | 1.27 | 1.14 | 1.23 | 1.12 | 1.13 |
| Boric acid | 1.15 | 1.17 | 1.17 | 1.24 | 1.08 | 1.22 | 1.22 |
| Nano boron | 1.32 | 1.02 | 1.19 | 1.14 | 1.28 | 1.26 | 1.25 |
| Mean | 1.24 | 1.10 | 1.21 | 1.17 | 1.20 | 1.20 | 1.20 |
| **Total Soluble Solids (Brix<sup>°</sup>)** |  |  |  |  |  |  |  |
| Control | 18.2 | 21.9 | 20.8 | 20.3 | 21.4 | 21.9 | 20.4 |
| Boric acid | 19.4 | 23.8 | 22.1 | 21.4 | 22.6 | 23.3 | 21.4 |
| Nano boron | 19.8 | 25.3 | 23.5 | 22.5 | 23.4 | 24.2 | 21.9 |
| Mean | 19.1 | 23.7 | 22.1 | 21.5 | 22.5 | 23.1 | 21.4 |
| **L.S.D. 0.05** |  |  |  |  |  |  |  |
|  | B | G | R | B x G | B x R | G x R | B x G x R |
| Sucrose | 0.11 | 0.65 | 0.30 | 0.49 | 0.51 | 0.60 | 0.05 |
| Purity | 0.08 | 0.70 | 0.25 | 0.25 | 0.80 | 0.42 | 0.10 |
| αAmino N | 0.10 | 0.18 | 0.09 | 0.11 | 0.21 | 0.09 | 0.08 |
| TSS | 0.25 | 1.15 | 0.35 | 0.40 | 0.75 | 0.58 | 0.15 |

**B = Boron sources; G = Plant growth regulator sources and R = Rate of Plant growth regulator**

Also, (Asil et al., 2011) who indicated drenching with gibberellin increased the floral stalk height as compared to the control. It may be attributed to the effect of gibberellin in stimulating and accelerating cell division, increasing cell elongation and enlargement, or both (Hartmann and Kester, 1963).

Regard to data in (Table 4) the foliar application of boric acid and B-NPs as a boron sources on sugar beet plants observed that nano boron source was more effective than boric acid for sucrose, purity, α- amino nitrogen and total soluble solids with values (16.48, 16.32 and 16.69, 16.41%), (80.47, 80.25 and 80.62, 80.31%), (1.15, 1.37 and 1.32, 1.21%) and (19.4, 18.4 and 19.8, 18.7 Brix□), respectively. These results were in agreement with (Naderi et al., 2011) who dedicated that application of nanofertilizers instead of common fertilizers, where nutrients are provided to plants gradually and in a controlled manner. Meanwhile, the nanotechnology increases the application efficiency of fertilizers, decreases pollution and risks of chemical fertilization.

Finally, the interaction analysis for 75% NPK from the recommended dose accompanied with foliar application of gibberellin at rate 100 mg l^−1^ and proline at rate 200 mg l^−1^ with nano boron was the most effective treatment than the same interaction with boric acid.

### 3.4. Bio-chemical components of sugar beet

The represented data in Table 5 showed that the foliar application of gibberellin at 100 mg l^−1^ was the most effective treatment followed by proline at 200 mg l^−1^ as in betaine content 0.172 and 0.171 g 100 g^−1^, choline content 8.40 and 7.93 mg 100g^−1^ FW and total carbohydrate content 14.8 and 14.3 g 100g^−1^). These results in harmony with (El-Sherbeny and Da Silva, 2013) who reported that proline proved to be successful agents in improving growth and yield characters of beet plants, especially at 100 mg·L^−1^ and 200 mg·L^−1^ also, GA_3_ increases the chlorophyll concentrations in leaves by increasing the numbers and sizes of chloroplasts and enhances the ultra-structural morphogenesis of plastids (Richard N. A., 1996). In general, photosynthetic efficiency increases along with the chlorophyll concentration. Thus, exogenous GA_3_ indirectly causes the increase in chlorophyll (Ashraf et al., 2002) that resulted in the accumulation of more dry mass (Khan, 1996). High concentration of endogenous GA_3_ by increasing the gibberellin concentration treatment (Wang et al., 2015). In addition, GA_3_ caused the increase in the endogenous Indole acetic acid contents (Reid and Davies, 1992). The previous studies have illustrated that the exogenous application of proline mitigated the decreased plant growth caused by stress is through increasing antioxidant system, relieving oxidative damage, improving the synthesis of compatible solutes, and accelerating proline accumulation, which reflected on enhancing photosynthesis (Szabados and Savouré, 2010); (Anjum et al., 2011). Also, Sun Physiologically, a major function of GAs in higher plants can be generalized as stimulating organ growth through enhancement of cell elongation and, in some cases, cell division. In addition, GAs promotes certain developmental switches, such as between seed dormancy and germination, juvenile and adult growth phases, and vegetative and reproductive development.

**Table 5.** Integrated effect of plant growth regulators and macro-nutrient fertilizer rates with different boron sources on biochemical contents

| Boron sources (B) | Gibberellin (mg l<sup>-1</sup>) |  |  |  | Proline (mg l<sup>-1</sup>) |  |  |
| --- | --- | --- | --- | --- | --- | --- | --- |
|  | Control | 100 | 200 | Mean | 100 | 200 | Mean |
| **Betaine (g 100 g<sup>-1</sup> root FW)** |  |  |  |  |  |  |  |
| Control | 0.162 | 0.172 | 0.160 | 0.162 | 0.160 | 0.171 | 0.161 |
| Boric acid | 0.168 | 0.194 | 0.166 | 0.174 | 0.174 | 0.180 | 0.172 |
| Nano boron | 0.173 | 0.208 | 0.172 | 0.185 | 0.187 | 0.198 | 0.187 |
| Mean | 0.168 | 0.191 | 0.166 | 0.174 | 0.174 | 0.183 | 0.173 |
| **Choline (mg 100 g<sup>-1</sup> root FW)** |  |  |  |  |  |  |  |
| Control | 7.45 | 8.40 | 7.75 | 7.81 | 7.78 | 7.93 | 7.72 |
| Boric acid | 7.51 | 8.68 | 7.91 | 7.95 | 7.81 | 8.15 | 7.74 |
| Nano boron | 7.58 | 8.82 | 7.90 | 8.10 | 7.95 | 8.32 | 7.84 |
| Mean | 7.51 | 8.63 | 7.98 | 7.92 | 7.85 | 8.14 | 7.80 |
| **Total Carbohydrate (g 100g<sup>-1</sup>)** |  |  |  |  |  |  |  |
| Control | 13.1 | 14.8 | 14.0 | 13.9 | 13.5 | 14.3 | 13.6 |
| Boric acid | 13.7 | 15.6 | 14.7 | 14.4 | 13.9 | 14.8 | 13.8 |
| Nano boron | 13.9 | 15.9 | 15.0 | 14.9 | 14.1 | 15.5 | 14.2 |
| Mean | 13.6 | 15.4 | 14.6 | 14.5 | 13.8 | 14.9 | 14.0 |
| **L.S.D. 0.05** |  |  |  |  |  |  |  |
|  | B | G | R | B x G | B x R | G x R | B x G x R |
| Betaine | 0.001 | 0.005 | 0.01 | 0.002 | 0.01 | 0.05 | 0.001 |
| Choline | 0.05 | 0.30 | 0.50 | 0.09 | 0.10 | 0.15 | 0.20 |
| Total Carb. | 0.10 | 1.00 | 0.75 | 0.25 | 0.11 | 0.31 | 0.90 |

**B = Boron sources; G = Plant growth regulator sources and R = Rate of Plant growth regulator**

Table 5 showed the foliar application of nano boron give higher results for betaine, choline and total carbohydrate than boric acid foliar application this may due to (Nilanjan, 2013) who stated that, one benefit of the boron nanofertilizer described in embodiments may be extremely low cost and high efficiency. Some embodiments described herein provide a highly effective means of nano-fertilization by administration of boric acid nanoparticles to plants. Nano scale boric acid released from the surface of metal nanoparticles of embodiments herein can be a highly efficient boron fertilizer. Other benefits of the boron coated metal nanoparticles described herein include increased boron content in plants resulting in increased chlorophyll content, number of leaves, total biomass, total yield, and lowered soluble and reducing sugars.

Finally, the interaction between the foliar application of gibberellin at 100 mg l^−1^ and proline at 200 mg l^−1^ with nano boron with 75% of recommended dose give the highest response for most parameter under study of sugar beet plant these results was in agreement with (Seadh and El-Metwally, 2015) who reported that adding of plant growth regulators with addition mineral fertilizers with 80% of recommended dose in order to maintain high productivity and quality of plant at the same time reduce production costs and environmental pollution under the environmental condition in Egypt.

## 4. Conclusion

From previously data obtained could be concluded that the using of plant growth regulators as a proline or gibberellin reduced the consumption of mineral fertilizers and possibility of replacing the gibberellin by proline.

The application of nanotechnology in agriculture as using nano boron (B-NPs) increased the application efficiency, decreased pollution and risk of fertilizers used and increased plant quality.

